# Neuroinflammation and Spinal Cord Pathology Associated with Hypertension in Spontaneously Hypertensive Stroke-Prone Rats

**DOI:** 10.1101/2024.10.12.618038

**Authors:** H. Noyan, B.H.J. Juurlink

**Affiliations:** Cameco MS and Neuroscience Research Centre; College of Medicine, University of Saskatchewan

## Abstract

Hypertension is a major contributor to cardiovascular and cerebrovascular disease, leading to significant central nervous system (CNS) damage, including in the spinal cord. This study examines the spinal cord in uncontrolled hypertension of stroke-prone spontaneously hypertensive rats (SHRsp) compared to normotensive Wistar-Kyoto rats (WKY). Histological analyses, including Sudan Black B staining, NADPH-diaphorase histochemistry, immunohistochemistry, and Western blotting, were used to assess inflammation, gliosis, oxidative stress, and axonal injury.

SHRsp rats displayed extensive spinal cord pathology, including macrophage infiltration, astrocyte hypertrophy, and demyelination. ED1 immunostaining revealed significant microglial activation in SHTsp, particularly in the ventrolateral white matter, while Sudan Black B staining showed edema and lipid-laden macrophages in dilated perivascular spaces. Increased NADPH-diaphorase activity in SHRsp indicated heightened nitric oxide synthase activity. Glial fibrillary acidic protein (GFAP) immunostaining confirmed significant astrogliosis, with hypertrophic astrocytes more prominent in SHRsp. Western blot analysis showed elevated amyloid precursor protein (APP) levels in SHRsp, associated with axonal injury and oxidative stress. APP-positive phagocytes in white matter further suggested active axonal degeneration.

These findings demonstrate significant neuroinflammation, gliosis, and spinal cord tissue damage in hypertensive rats, highlighting the spinal cord’s vulnerability to hypertension. This underscores the importance of addressing hypertensive damage in the CNS, particularly in patients at risk of neurodegenerative or motor impairments.

## INTRODUCTION

Hypertension is a significant, preventable risk factor for cardiovascular and cerebrovascular mortality and morbidity worldwide ^1^. Despite the widespread availability of pharmacological treatments, the global control rates of this disease remained alarmingly low ^2^. In many cases, essential hypertension remains asymptomatic until it reaches an advanced stage ^3^. Unfortunately, this delayed diagnosis increases the risk of stroke, myocardial infarction, congestive heart failure and renal failure in affected individuals ^4^.

The brain and arterial blood vessels are primary targets of hypertensive damage in patients. In fact, chronic hypertension has been linked to conditions such as lacunar strokes ^5^, dementia ^2^, and cognitive impairments ^6^. Additionally, hypertension is associated with an increased prevalence of white matter lesions, especially in individuals with poorly controlled hypertension, even those under treatment ^7^.

Spontaneously hypertensive rats (SHR) and their stroke-prone counterparts (SHRsp) have long been established as valuable animal models for studying essential hypertension ^8^. Research on these models has revealed various pathological changes in the brain regions of SHR and SHRSPsp, including cerebral atrophy ^9^, blood-brain barrier dysfunction ^10^, vascular abnormalities ^11^, neuroinflammation ^12^, neuronal loss, astrogliosis ^13^, and white matter lesions ^12, 14, 15^. However, despite these extensive studies on the brain, the effects of hypertension on the spinal cord - a key region of the central nervous system (CNS) responsible for essential motor and autonomic functions - have been largely overlooked. In our previous investigation, which focused on dietary interventions to mitigate CNS inflammation in 10-month-old SHRsp, we observed significant pathological changes in the spinal cord ^16^. This unexpected finding suggests that the spinal cord, like the brain, may be vulnerable to hypertensive damage.

Given the spinal cord’s critical role in maintaining homeostasis and regulating physiological processes, it is important to address this gap in the research. Understanding the histopathological changes in the spinal cord due to undiagnosed/uncontrolled hypertension could reveal new therapeutic targets and improve the management of CNS damage in hypertensive patients, especially those exhibiting neurodegenerative or motor impairments. We hypothesize that uncontrolled hypertension induces pathological changes in the spinal cord similar to those seen in other CNS regions, including neuroinflammation, gliosis, and tissue degeneration. The aim of this descriptive study is to systematically investigate and characterize the histopathological effects of uncontrolled hypertension on the spinal cord in young SHRSP, contributing to a broader understanding of the CNS consequences of this condition.

## MATERIALS AND METHODS

### Animal Groups

Male stroke-prone spontaneously hypertensive rats (SHRsp) and age-matched Wistar-Kyoto rats (WKY) were housed under standard conditions of light/dark cycles and controlled humidity. The animals were fed Purina rat chow and given water ad libitum. All procedures involving the animals were conducted in accordance with the Canadian Council of Animal Care Guidelines. For this study, five animals from each group (N=5/group) were assessed for pathological changes in the spinal cord at 40 weeks of age.

### Tissue Preparation

Rats were deeply anesthetized with halothane and transcardially perfused via the ascending aorta with heparinized 0.05 M phosphate-buffered saline (PBS), pH 7.4. In a subset of animals, perfusion was continued with 4% paraformaldehyde in PBS. After perfusion, the spinal cords were carefully dissected and immersed in the same fixative overnight at 4°C. For cryoprotection, the tissues were incubated in a 30% sucrose solution for three days at 4°C. Spinal cords were then embedded in OCT compound (Somagen Diagnostics, Inc., Edmonton, AB, Canada), and cross-sections (10-14 µm thick) were prepared. The sections were mounted on Superfrost Plus-coated slides (VWR) and stored at -70°C until further analysis.

### Staining Methods

#### A. Sudan Black B Staining

To assess potential myelin degradation and depletion, we employed a modified Sudan Black B staining technique ^17^. The sections were counterstained with 0.02% neutral red in acetate buffer (pH 3.3), mounted in glycerin jelly, and coverslipped for imaging.

#### B. Enzyme Histochemistry

NADPH-diaphorase staining was performed on thoracic spinal cord sections using a reaction mixture containing 1 mg/ml β-NADPH (Sigma), 0.5 mg/ml Nitro Blue Tetrazolium (NBT, Sigma), and 0.3% Triton X-100 in PBS, incubated at 37°C in the dark ^18^. Controls for the reaction were prepared by omitting NADPH, resulting in the absence of staining except for a reddish monoformazan background. This technique was also used for double staining with GFAP immunohistochemistry or with an acidic neutral red counterstain.

#### C. Immunohistochemistry

Sections were air-dried for 10 minutes and rehydrated in 0.1 M PBS. To block endogenous peroxidase activity, sections were incubated with 0.3% H2O2 in methanol for 30 minutes and washed thoroughly with PBS. To reduce non-specific staining, sections were preincubated in 10% normal horse serum (Sigma) in PBS for 1 hour at room temperature. The sections were then incubated overnight at 4°C with primary antibodies, including: mouse monoclonal ED1 (1:1000, Serotec, UK) to detect activated macrophages/microglia, anti-GFAP (1:400, GA-4, Sigma), anti-nitrotyrosine (1:300, rabbit polyclonal, Upstate Biotechnology, NY, USA), Goat antirabbit anti-ATF3 (1:300, sc-188, Santa Cruz Biotech. Inc.), or anti-amyloid precursor protein (APP, 1:1000, Sigma). Following three PBS washes, sections were incubated with biotin-labeled secondary antibodies (1:1000, Vector Laboratories, Burlington, ON, Canada) or Cy3 Goat anti-Rabbit IgG (For ATF3) for 1 hour at room temperature before mounting and fluorescence microscopy. After incubation with the avidin-biotin complex (ABC, Vector), immunoreactivity was visualized using 3,3′-diaminobenzidine (DAB) with 0.03% H2O2 as the chromogen. For some sections, a nickel solution was applied to intensify the signal. After several washes, sections were dehydrated in ethanol, cleared with xylene, and coverslipped using Entellan (VWR). Negative controls were conducted by omitting either the primary or secondary antibody. The percent object area of GFAP-positive profiles in the white matter of the thoracic spinal cord in both WKY and SHRSP rats was quantified using Northern Eclipse software.

### Microscopy

All images were captured using a Zeiss Axioskop microscope equipped with a Sony DXC-950 camera.

### Western Blotting

Spinal cord samples were prepared for Western blot analysis to measure APP abundance, as previously described ^16^. Rabbit polyclonal anti-GAPDH (Santa Cruz) was used as a loading control. HRP-conjugated secondary antibodies were visualized using chemiluminescence. Blots were scanned, and band optical densities were analyzed using ImageJ software (NIH). The ratio of APP to GAPDH was calculated.

### Statistical Analysis

Data for GFAP percent object area and Western blot analyses are presented as mean ± SE. An unpaired Student’s t-test was used for group comparisons, with significance set at p<0.05. GraphPad Prism v5.0 (GraphPad Software, Inc., La Jolla, CA, USA) was used for statistical analysis.

## RESULTS

A significant inflammatory and edematous state was observed in the spinal cord of 40-week-old SHRsp. The study revealed selective infiltration of macrophages in the white matter, astroglial hypertrophy and activation in both white and gray matter, increased expression of nitrotyrosine and amyloid precursor protein (APP), and the presence of inducible nitric oxide synthase (iNOS) in neurons and non-neuronal cells in thoracolumbar spinal cord sections of SHRsp compared to WKY.

Immunostaining with the anti-ED1 antibody revealed no positive cells in the spinal cord of WKY (Fig. 1A), but massive macrophage/microglial activation in the white matter of SHRsp (Figs. 1B-E). ED1-positive cells were ovoid, ranging from 10 to 15 micrometers in diameter (sometimes larger), and contained abundant lysosomal vesicles, indicative of active phagocytosis. The distribution of ED1 immunoreactivity was not uniform within the white matter, with higher numbers of ED1-positive cells located in the outer portions of the ventral white matter, extending from the dorsolateral to medioventral regions (Figs. 1B-E). The dorsal funiculus (Fig. 1C) exhibited fewer ED1-positive cells than the ventrolateral white matter (Figs. 1D-E). These phagocytes were found individually or in small groups near myelinated fibers, around small blood vessels, and in dilated perivascular or intraparenchymal spaces. Inflammation in the SHRsp spinal cord was also associated with increased protein nitrosylation (Fig. 1G) compared to age-matched WKY (Fig. 1F).

**Figure 1.**
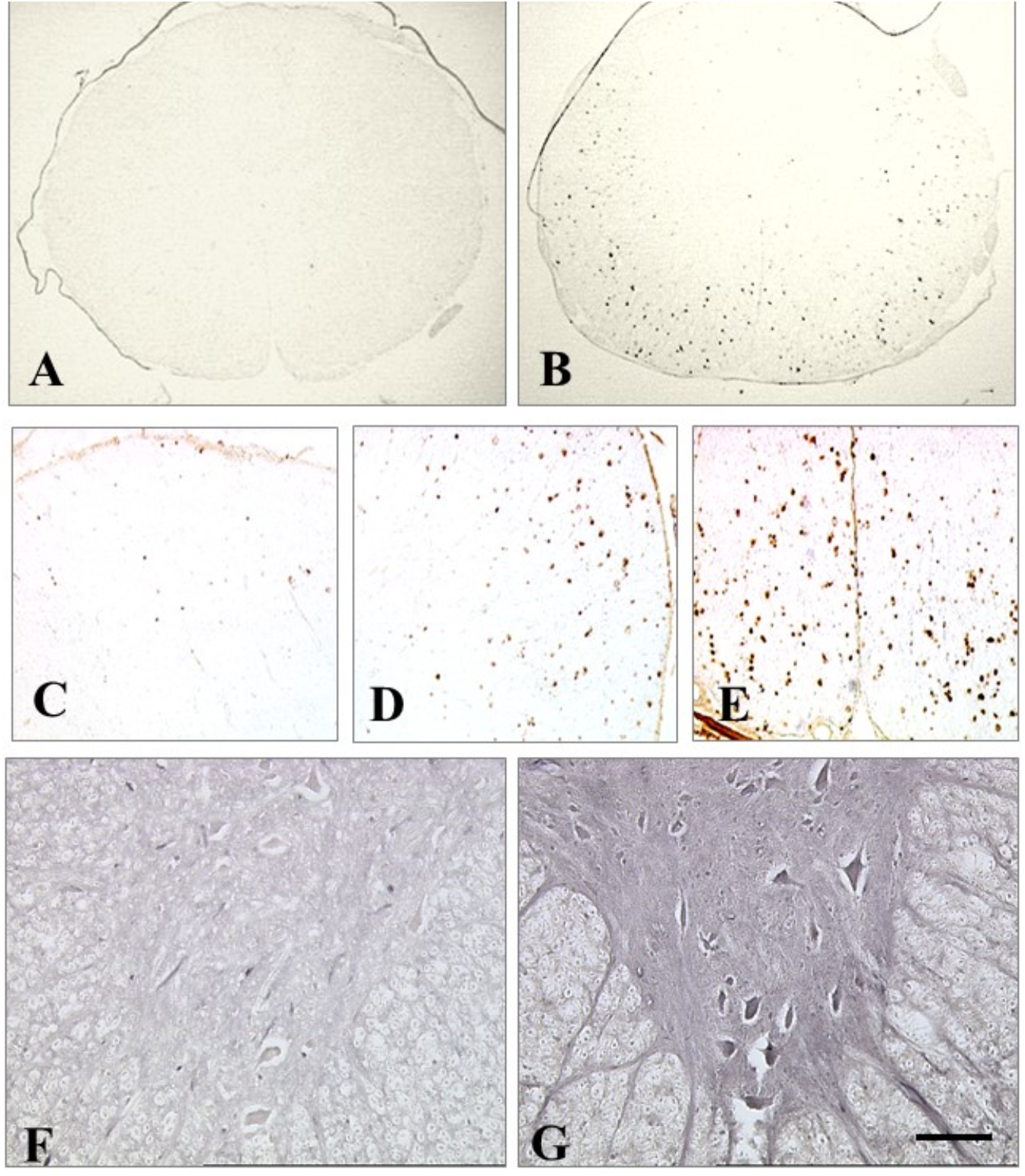
Representative low magnification photographs demonstrating lack of ED1-positive cells in the thoracic spinal cord (TSC) tissue of WKY rats (A) while white matter of the TSC in SHRsp is infiltrated with many ED1-positive cells (B). ED1-positive cells are present in the dorsal (C), latera (D) and ventral (E) columns of the spinal cord in the SHRsp. Staining with anti-nitrotyrosine antibody showed weak immunoreactivity in the gray- and white matter of the TSC tissue of the WKY (F) but was quite significant in the SHRsp (G). Scale bar: 700 micrometers for A and B, 100 micrometers for C-E, 50 micrometers for F & G.

Sudan Black B staining of the WKY spinal cord revealed normal white matter architecture. In contrast, SHRsp spinal cords exhibited dilated perivascular and intraparenchymal spaces in the white matter, suggesting edema (Fig. 2). These spaces were frequently occupied by large, lipid-containing macrophage-like cells (Fig. 2D). Phagocytes containing sudanophilic granules, indicative of myelinophagia, were observed around the dilated spaces (Fig. 2E) and were distributed throughout the parenchyma (Fig. 2F).

**Figure 2.**
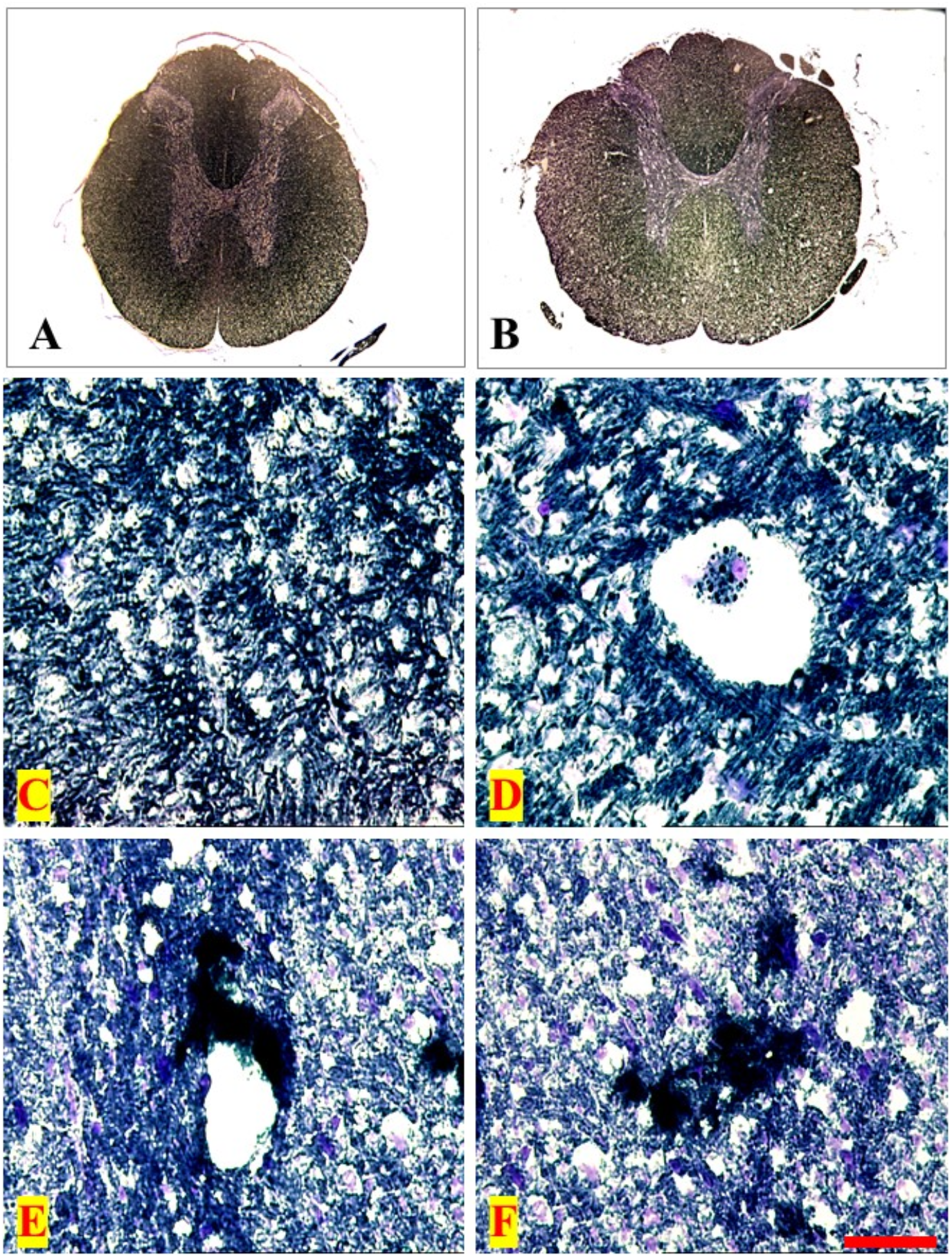
Sudan black B staining. Overview of the thoracic spinal cord (TSC) of WKY (A) and SHRsp (B), shows edema and intraparenchymal dilatations. C, shows normal TSC tissue in the WKY while in the SHRsp dilated intraparenchymal spaces in the white matter (D) are occupied with sudanophilic macrophages. E and F show the patchy accumulation of these macrophages in the white matter of the SHRsp. Overall evidence for demyelination are shown in D-F. Scale bar: 1000 micrometers for A&B; 70 micrometers for C&D; and 50 micrometers for E&F.

The NADPH-diaphorase reaction product was localized in various regions, including: (1) the superficial laminae (laminae I-IV) of the dorsal horn gray matter, with higher activity in SHRsp than WKY; (2) microvasculature, which exhibited less staining in WKY than SHRsp; (3) a few fusiform and ovoid cells in both WKY and SHRsp; (4) large, stellate motoneurons in the ventral horn gray matter in SHRsp; and (5) cells with astrocyte morphology in SHRsp. Double-staining with GFAP confirmed the identity of these astrocytes. Two major differences were noted between the spinal cord sections of SHRsp and WKY: (a) a marked upregulation of diaphorase activity in the dorsal horn gray matter (laminae I-IV) of SHRsp, evidenced by an increase in diaphorase-positive punctate elements in the neuropil (Figs. 3A & B); and (b) increased NADPH-diaphorase activity in the vasculature of SHRsp, particularly in endothelial and possibly perivascular cells in the walls of blood vessels penetrating the white matter (Figs. 3C & D). Since NADPH-diaphorase staining marks nitric oxide synthase activity, it reflects the activity of all three isoforms (eNOS, nNOS, and iNOS). Previous studies indicated that iNOS accounts for part of this expression in the SHRsp spinal cord ^16^.

**Figure 3.**
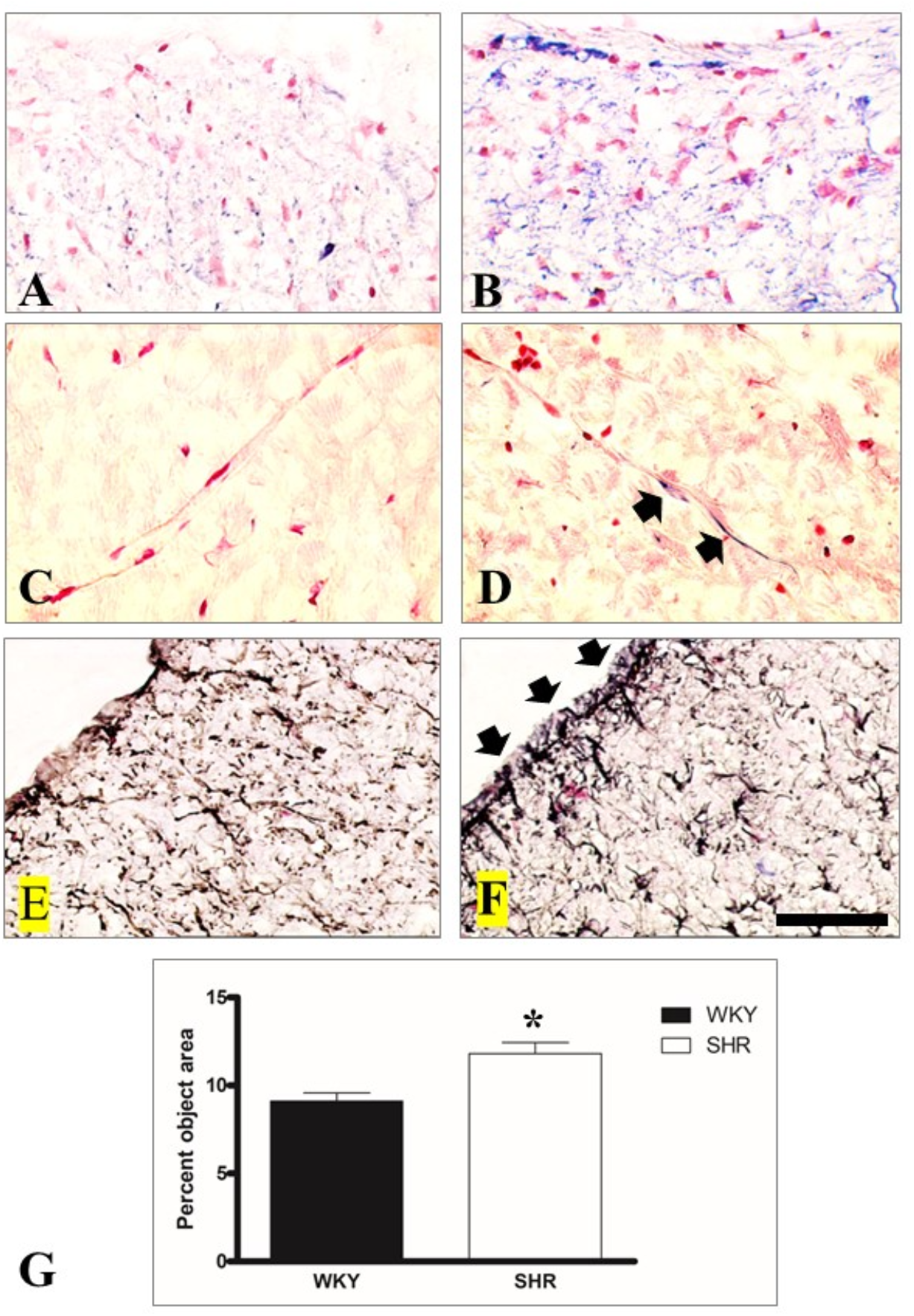
NADPH-diaphorase staining of the dorsal horn of TSC compared in WKY(A) and SHRsp (B). Neutral red counterstaining is used to identify the nuclei and staining the background. A considerable increase in diaphorase staining is visible in SHR. Pictures C and D, present increased diaphorase activity in perivascular and parenchymal tissues in SHRsp (D) compared to WKY(C). Double-staining for NADPH-d and GFAP also showed astrogliosis in the superficial part of the dorsal Horn in SHRsp (F) compared to WKY(E). Measurement of percent object area (POA) of GFAP-positive astrocytes in the white matter showed increased POA in SHRsp compared with WKY (G). Scale Bar: 50 micrometers.

GFAP-immunoreactive process-bearing astrocytes were present throughout the gray and white matter of both WKY (Figs. 3E & 4A) and SHRsp (Figs. 3F & 4B). However, staining was more intense, and astrocytes appeared larger in SHRsp, indicating astroglial hypertrophy and activation. This activation was particularly prominent around the central canal and the superficial laminae, including the zone of Lissauer in the dorsal horn (Figs. 3E & F). In the white matter, activated astroglia in SHRsp displayed hypertrophy and strong GFAP immunoreactivity. Quantification of GFAP-positive profiles in the white matter of the thoracic spinal cord revealed a significant increase (P<0.05) in astrocyte coverage in SHRsp compared to WKY (Fig. 3G). Additionally, cells resembling endothelial or perivascular cells showed increased diaphorase activity in SHRsp (Figs. 3C & D).

Neurons in the intermediolateral nucleus (IML) of the thoracic spinal cord were identified based on their location and NADPH-diaphorase staining (Anderson CR, Neurosci Lett, 1992, Figs. 4A & B). In SHRsp, these neurons were surrounded by hypertrophic astrocytes (Figs. 4A & B) and expressed iNOS (Figs. 4C & D).

**Figure 4.**
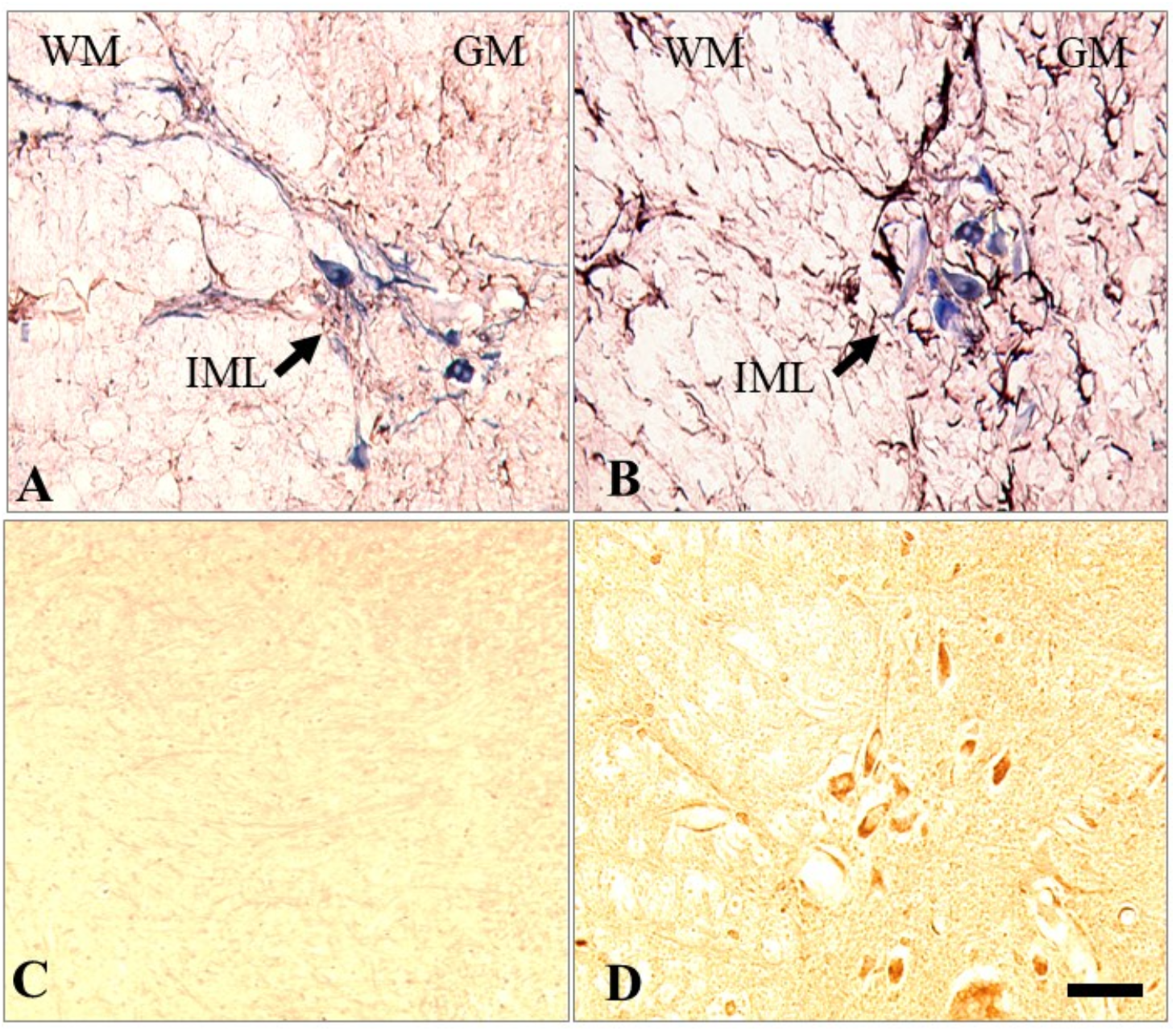
Double-staining for NADPH-d and GFAP showed astrogliosis in and around intermediolateral horn (IML) of the SHRsp (B) compared to WKY (A). IML neurons in SHRsp express iNOS (D) while WKY rats do not express this inducible enzyme (C). Scale Bar: 200 micrometers.

APP-immunoreactivity was diffusely present in the somas and primary dendritic segments of neurons and glial cells in WKY (Figs. 5A & C). In SHRsp, neuronal somas, processes, and glial cells exhibited much stronger APP staining (Figs. 5B & D). Macrophage/microglia-like cells in SHRsp also showed intense APP-immunoreactivity (Figs. 5E & F). Axonal injury was evident in SHRsp, as shown by patchy and diffuse intra-axonal APP accumulations (Fig. 5F). Furthermore, APP accumulations were detected in phagocytes and groups of cells in the white matter of SHRsp, but not in WKY (Figs. 5D-F). Increased APP expression in the spinal cord of SHRsp was also confirmed by Western blot analysis (Fig. 6).

**Figure 5.**
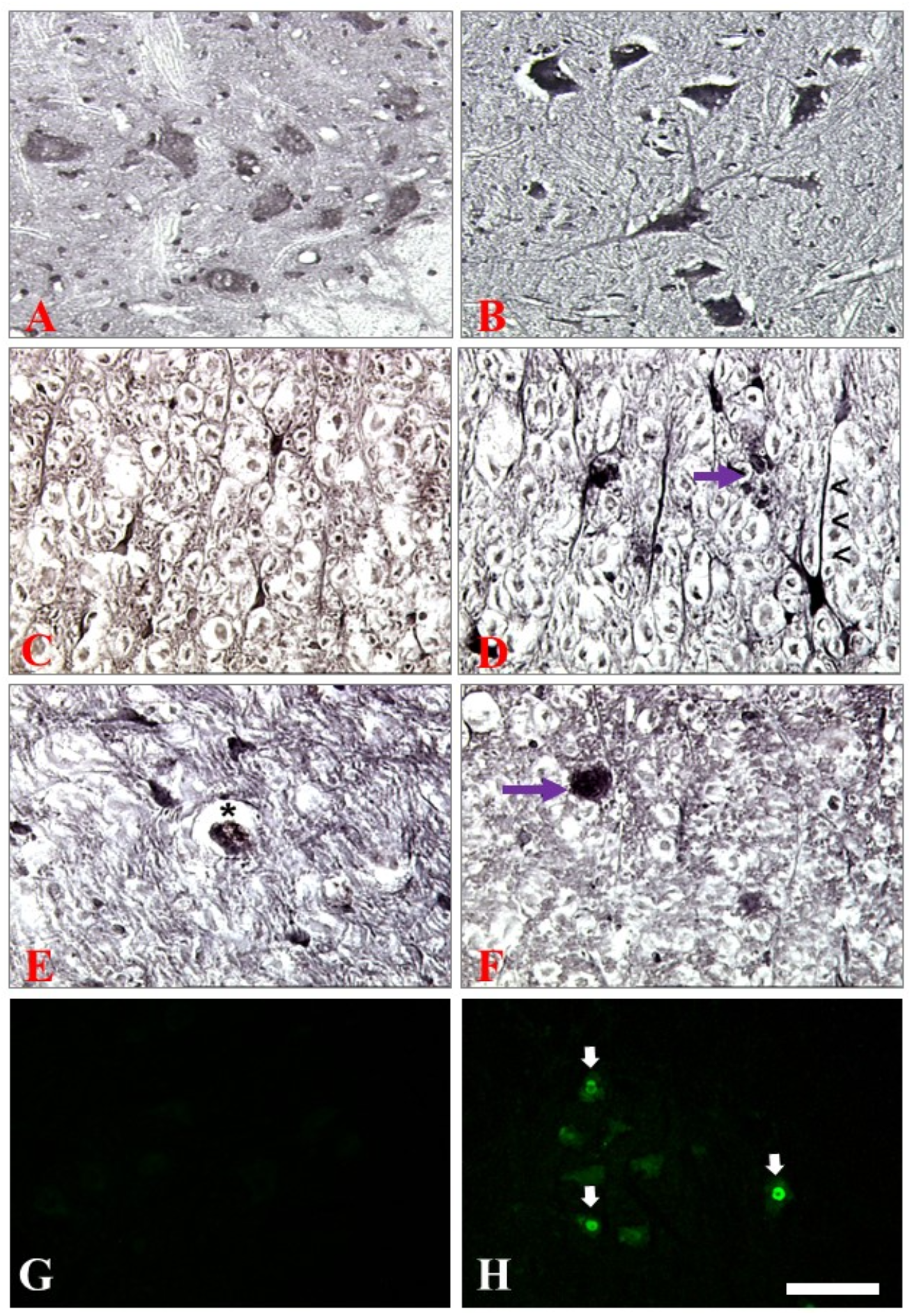
Pattern of APP immunoreactivity is compared in spinal grey matter (A&B) and white matter (C&D) in TSC of WKY and SHRsp. APP-immunoreactivity is limited to the cell bodies and proximal processes of the neuronal and glial cells in WKY (A&C) while in SHRsp (B&D) motoneurons and hypertrophic astrocytes show APP-immunoreactivity both in cell body and further in distal part of their dendritic arborizations. Strong APP-immunoreactivity in the hypertrophic astrocytes (D) and their processes(arrowheads) and also in the phagocytes. Phagocytes expressing APP-immunoreactivity are present in the spinal white matter of the SHRsp (D, arrow & E, asterisk, F, arrow). Some neuronal nuclei are labelled express ATF3 in SHRsp’s ventral horn (H), and not WKY (G). Scale bar: 150 micrometers for A and B, 50 micrometers for C-H.

**Figure 6.**
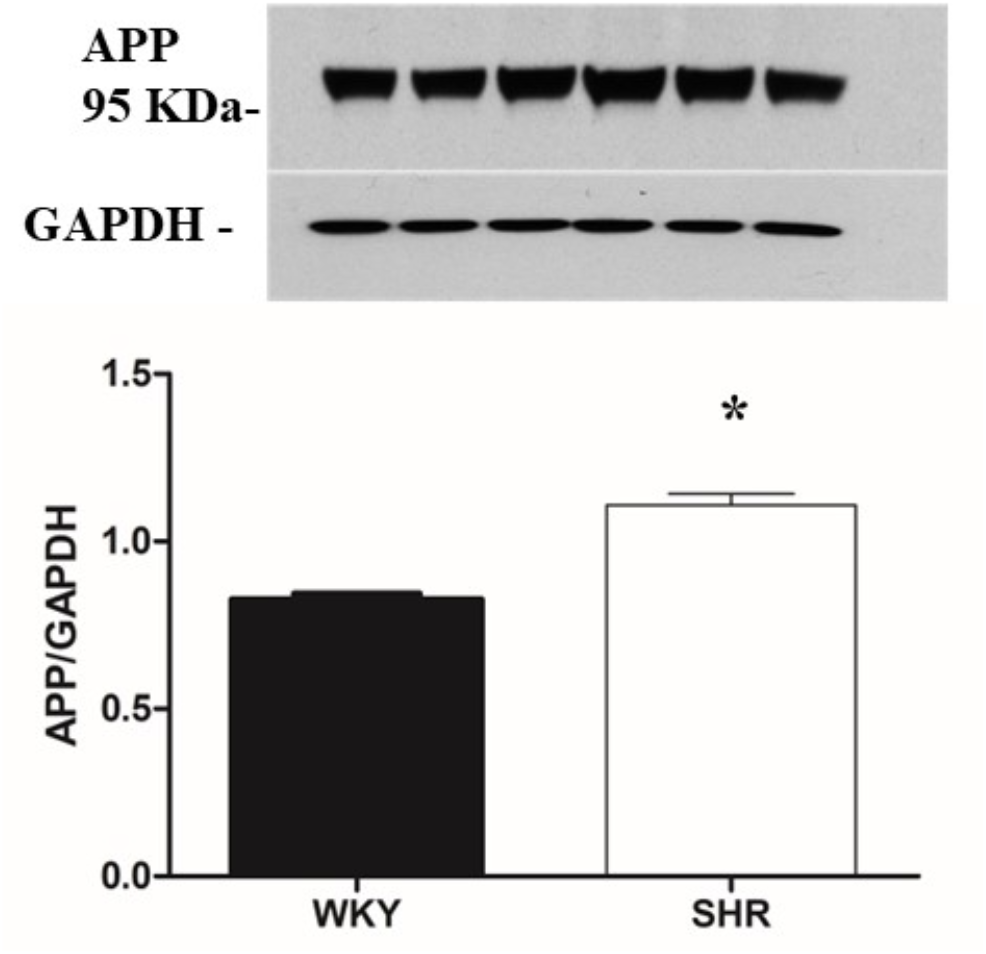
Representative Western blot analysis of APP protein abundance in the thoracic spinal cord extracts of WKY compared to SHRsp using GAPDH as loading control. APP protein expression levels is increased in SHRsp. Data shown are means ±SE; \**P* < 0.05.

## DISCUSSION

The present study demonstrates that uncontrolled hypertension in 40-week-old SHRsp rats is associated with significant histopathological changes in the spinal cord, particularly in the white matter. Notably, we observed extensive macrophage infiltration, astroglial activation, and increased expression of key markers associated with oxidative stress and neuroinflammation, such as nitrotyrosine, APP, and iNOS. These changes included a large number of ED1-positive cells, indicative of activated monocytes and microglia, as well as myelin depletion and increased NADPH-diaphorase activity in both gray and white matter. The glial activation and white matter lesions were most pronounced in the outer portions of the ventral and lateral funiculi, areas critical for transmitting nociceptive and proprioceptive sensory information. These findings align with similar observations in the hypertensive brain, but importantly, they extend our understanding of how hypertension affects the spinal cord, highlighting its vulnerability to neuroinflammatory and degenerative processes.

Hypertension is a significant risk factor for ischemic heart disease, stroke, cardiovascular diseases, chronic kidney disease, and dementia ^2^. SHR, a well-established model for essential (neurogenic) hypertension ^19^, is characterized by neuroinflammation and elevated sympathetic nerve activity ^20, 21^. In this study, we used the anti-ED1 antibody, which recognizes rat lysosomal membrane antigen in activated macrophages and microglia ^22, 23^. ED1 immunohistochemistry revealed significant microgliosis in the inflamed white matter of the SHRsp spinal cord, consistent with previous studies that have reported similar changes in the white matter of the brain in SHR and SHRsp ^24, 25^. Interestingly, microglial involvement in the regulation of sympathetic outflow ^26, 27^ suggests that microglial activation in the SHRsp spinal cord may also contribute to sympathetic dysregulation observed in hypertension.

Our findings suggest that the SHRsp model exhibits not only hypertensive damage but also features of accelerated aging. Microgliosis in the SHRsp spinal cord was similar to that observed in aged rat models, such as 30-month-old Sprague-Dawley rats ^28^. This suggests that hypertension may accelerate age-related pathological processes, particularly in the central nervous system (CNS), making SHRsp a useful model for studying both hypertension and aging-related neurodegeneration.

Oxidative stress is a major factor in the pathogenesis of hypertension ^29^, and nitrotyrosine, a biomarker of oxidative and nitrative stress, was found in the spinal cord tissues of SHRsp in this study. This biomarker has been implicated in chronic CNS inflammation and demyelination ^30, 31^. Sympathetic dysregulation, a hallmark of essential hypertension, is also believed to be maintained by elevated sympathetic nerve activity ^32, 33^. In this context, our study identified astrogliosis and iNOS induction in preganglionic sympathetic neurons of the intermediolateral (IML) region of the SHRSP thoracolumbar spinal cord. The IML region plays a key role in cardiovascular regulation and nociceptive processes ^34, 35^. The cytological changes observed in this region may be of functional and clinical importance, highlighting the complex interplay between neuroinflammation, oxidative stress, and hypertension-induced damage.

A striking feature of this study was the elevated amyloid precursor protein (APP) expression in both the gray and white matter of the SHRsp spinal cord. APP, a transmembrane protein implicated in neural plasticity, memory, and neuroprotection ^36, 37^, was strongly expressed in neurons, astrocyte-like cells, and activated microglia/macrophages in the hypertensive rats. The presence of APP in neurons and glial cells has been documented in both intact rats and mice ^38, 39^, and it is known to regulate immune cell functions ^40^. In SHRsp, the increased APP expression may reflect a proinflammatory state in the CNS, potentially contributing to neuronal damage and impaired microglial function ^41, 42^. Moreover, we observed plaque-like accumulations of APP in the spinal white matter, suggestive of axonal injury, which could disrupt axonal transport and neural transmission. This axonal damage was further confirmed by ATF-3 staining, a marker of neuronal injury, in motor neurons of the ventral horn ^43^.

It is important to recognize that not all CNS changes in hypertensive rats are strictly dependent on elevated blood pressure. For instance, both hypoalgesic and hyperalgesic responses have been reported in SHRsp, independent of blood pressure ^44^. Similarly, SHRsp dorsal columns exhibit hyper-responsiveness to stimuli, even when the animals are normotensive from a young age ^45^. These functional alterations suggest that neuroinflammation and other histopathological changes in the spinal cord may contribute to sensory dysfunctions beyond the effects of hypertension alone.

In conclusion, our findings demonstrate that the spinal cord is a valuable model for investigating the histopathological and inflammatory changes associated with chronic hypertension. The SHRsp model highlights the contributions of oxidative stress, inflammation, and neurodegeneration, particularly in white matter and the preganglionic sympathetic neurons of the IML region. The observed axonal injury, demyelination, and neuroinflammatory responses are key features of hypertension-induced CNS pathology. Moreover, the accelerated aging-like changes in SHRsp further underscore the utility of this model in studying neurodegenerative processes. Future studies should explore the functional consequences of these histopathological changes and their potential reversibility through therapeutic interventions.

## Notes

### Competing Interest Statement

The authors have declared no competing interest.

## REFERENCES

1. Mills KT, Stefanescu A and He J. The global epidemiology of hypertension. Nat Rev Nephrol. 2020;16:223–237.

2. Zhou B, Perel P, Mensah GA and Ezzati M. Global epidemiology, health burden and effective interventions for elevated blood pressure and hypertension. Nat Rev Cardiol. 2021;18:785–802.

3. Ma J and Chen X. Advances in pathogenesis and treatment of essential hypertension. Front Cardiovasc Med. 2022;9:1003852.

4. Lackland DT and Weber MA. Global burden of cardiovascular disease and stroke: hypertension at the core. Can J Cardiol. 2015;31:569–71.

5. Pinto A, Tuttolomondo A, Di Raimondo D, Fernandez P and Licata G. Cerebrovascular risk factors and clinical classification of strokes. Semin Vasc Med. 2004;4:287–303.

6. Ungvari Z, Toth P, Tarantini S, Prodan CI, Sorond F, Merkely B and Csiszar A. Hypertension-induced cognitive impairment: from pathophysiology to public health. Nat Rev Nephrol. 2021;17:639–654.

7. Liao D, Cooper L, Cai J, Toole JF, Bryan NR, Hutchinson RG and Tyroler HA. Presence and severity of cerebral white matter lesions and hypertension, its treatment, and its control. The ARIC Study. Atherosclerosis Risk in Communities Study. Stroke. 1996;27:2262–70.

8. Trippodo NC and Frohlich ED. Similarities of genetic (spontaneous) hypertension. Man and rat. Circ Res. 1981;48:309–19.

9. Tajima A, Hans FJ, Livingstone D, Wei L, Finnegan W, DeMaro J and Fenstermacher J. Smaller local brain volumes and cerebral atrophy in spontaneously hypertensive rats. Hypertension. 1993;21:105–11.

10. Yang Y, Kimura-Ohba S, Thompson JF, Salayandia VM, Cosse M, Raz L, Jalal FY and Rosenberg GA. Vascular tight junction disruption and angiogenesis in spontaneously hypertensive rat with neuroinflammatory white matter injury. Neurobiol Dis. 2018;114:95–110.

11. Kaiser D, Weise G, Moller K, Scheibe J, Posel C, Baasch S, Gawlitza M, Lobsien D, Diederich K, Minnerup J, Kranz A, Boltze J and Wagner DC. Spontaneous white matter damage, cognitive decline and neuroinflammation in middle-aged hypertensive rats: an animal model of early-stage cerebral small vessel disease. Acta Neuropathol Commun. 2014;2:169.

12. Lin JX, Tomimoto H, Akiguchi I, Wakita H, Shibasaki H and Horie R. White matter lesions and alteration of vascular cell composition in the brain of spontaneously hypertensive rats. Neuroreport. 2001;12:1835–9.

13. Sabbatini M, Strocchi P, Vitaioli L and Amenta F. The hippocampus in spontaneously hypertensive rats: a quantitative microanatomical study. Neuroscience. 2000;100:251–8.

14. Masumura M, Hata R, Nagai Y and Sawada T. Oligodendroglial cell death with DNA fragmentation in the white matter under chronic cerebral hypoperfusion: comparison between normotensive and spontaneously hypertensive rats. Neurosci Res. 2001;39:401–12.

15. Sabbatini M, Baldoni E, Cadoni A, Vitaioli L, Zicca A and Amenta F. Forebrain white matter in spontaneously hypertensive rats: a quantitative image analysis study. Neurosci Lett. 1999;265:5–8.

16. Noyan-Ashraf MH, Sadeghinejad Z and Juurlink BH. Dietary approach to decrease aging-related CNS inflammation. Nutr Neurosci. 2005;8:101–10.

17. Gerrits PO, Brekelmans-Bartels M, Mast L, Gravenmade EJ, Horobin RW and Holstege G. Staining myelin and myelin-like degradation products in the spinal cords of chronic experimental allergic encephalomyelitis (Cr-EAE) rats using Sudan black B staining of glycol methacrylate-embedded material. J Neurosci Methods. 1992;45:99–105.

18. Usunoff KG, Kharazia VN, Valtschanoff JG, Schmidt HH and Weinberg RJ. Nitric oxide synthase-containing projections to the ventrobasal thalamus in the rat. Anat Embryol (Berl). 1999;200:265–81.

19. Zubcevic J, Jun JY, Kim S, Perez PD, Afzal A, Shan Z, Li W, Santisteban MM, Yuan W, Febo M, Mocco J, Feng Y, Scott E, Baekey DM and Raizada MK. Altered inflammatory response is associated with an impaired autonomic input to the bone marrow in the spontaneously hypertensive rat. Hypertension. 2014;63:542–50.

20. Haspula D and Clark MA. Neuroinflammation and sympathetic overactivity: Mechanisms and implications in hypertension. Auton Neurosci. 2018;210:10–17.

21. Oparil S, Acelajado MC, Bakris GL, Berlowitz DR, Cifkova R, Dominiczak AF, Grassi G, Jordan J, Poulter NR, Rodgers A and Whelton PK. Hypertension. Nat Rev Dis Primers. 2018;4:18014.

22. Damoiseaux JG, Dopp EA, Calame W, Chao D, MacPherson GG and Dijkstra CD. Rat macrophage lysosomal membrane antigen recognized by monoclonal antibody ED1. Immunology. 1994;83:140–7.

23. Suzuki H, Sugimura Y, Iwama S, Suzuki H, Nobuaki O, Nagasaki H, Arima H, Sawada M and Oiso Y. Minocycline prevents osmotic demyelination syndrome by inhibiting the activation of microglia. J Am Soc Nephrol. 2010;21:2090–8.

24. Amenta F, Strocchi P and Sabbatini M. Vascular and neuronal hypertensive brain damage: protective effect of treatment with nicardipine. J Hypertens Suppl. 1996;14:S29–35.

25. Kimura S, Saito H, Minami M, Togashi H, Nakamura N, Ueno K, Shimamura K, Nemoto M and Parvez H. Docosahexaenoic acid attenuated hypertension and vascular dementia in stroke-prone spontaneously hypertensive rats. Neurotoxicol Teratol. 2002;24:683–93.

26. Bi Q, Wang C, Cheng G, Chen N, Wei B, Liu X, Li L, Lu C, He J, Weng Y, Yin C, Lin Y, Wan S, Zhao L, Xu J, Wang Y, Gu Y, Shen XZ and Shi P. Microglia-derived PDGFB promotes neuronal potassium currents to suppress basal sympathetic tonicity and limit hypertension. Immunity. 2022;55:1466–1482 e9.

27. Li Y, Wei B, Liu X, Shen XZ and Shi P. Microglia, autonomic nervous system, immunity and hypertension: Is there a link? Pharmacol Res. 2020;155:104451.

28. Kullberg S, Aldskogius H and Ulfhake B. Microglial activation, emergence of ED1-expressing cells and clusterin upregulation in the aging rat CNS, with special reference to the spinal cord. Brain Res. 2001;899:169–86.

29. Loperena R and Harrison DG. Oxidative Stress and Hypertensive Diseases. Med Clin North Am. 2017;101:169–193.

30. Smith ME. Phagocytosis of myelin in demyelinative disease: a review. Neurochem Res. 1999;24:261–8.

31. van der Veen RC, Hinton DR, Incardonna F and Hofman FM. Extensive peroxynitrite activity during progressive stages of central nervous system inflammation. J Neuroimmunol. 1997;77:1–7.

32. Abboud FM. The sympathetic system in hypertension. State-of-the-art review. Hypertension. 1982;4:208–25.

33. Judy WV, Watanabe AM, Henry DP, Besch HR, Jr., Murphy WR and Hockel GM. Sympathetic nerve activity: role in regulation of blood pressure in the spontaenously hypertensive rat. Circ Res. 1976;38:21–9.

34. Anderson CR, Edwards SL, Furness JB, Bredt DS and Snyder SH. The distribution of nitric oxide synthase-containing autonomic preganglionic terminals in the rat. Brain Res. 1993;614:78–85.

35. Chowdhary S and Townend JN. Role of nitric oxide in the regulation of cardiovascular autonomic control. Clin Sci (Lond). 1999;97:5–17.

36. Cho Y, Bae HG, Okun E, Arumugam TV and Jo DG. Physiology and pharmacology of amyloid precursor protein. Pharmacol Ther. 2022;235:108122.

37. Muller UC, Deller T and Korte M. Not just amyloid: physiological functions of the amyloid precursor protein family. Nat Rev Neurosci. 2017;18:281–298.

38. Card JP, Meade RP and Davis LG. Immunocytochemical localization of the precursor protein for beta-amyloid in the rat central nervous system. Neuron. 1988;1:835–46.

39. Kawarabayashi T, Shoji M, Harigaya Y, Yamaguchi H and Hirai S. Amyloid beta/A4 protein precursor is widely distributed in both the central and peripheral nervous systems of the mouse. Brain Res. 1991;552:1–7.

40. Monning U, Konig G, Banati RB, Mechler H, Czech C, Gehrmann J, Schreiter-Gasser U, Masters CL and Beyreuther K. Alzheimer beta A4-amyloid protein precursor in immunocompetent cells. J Biol Chem. 1992;267:23950–6.

41. Blasko I, Stampfer-Kountchev M, Robatscher P, Veerhuis R, Eikelenboom P and Grubeck-Loebenstein B. How chronic inflammation can affect the brain and support the development of Alzheimer’s disease in old age: the role of microglia and astrocytes. Aging Cell. 2004;3:169–76.

42. von Bernhardi R, Ramirez G, Toro R and Eugenin J. Pro-inflammatory conditions promote neuronal damage mediated by Amyloid Precursor Protein and decrease its phagocytosis and degradation by microglial cells in culture. Neurobiol Dis. 2007;26:153–64.

43. Tsujino H, Kondo E, Fukuoka T, Dai Y, Tokunaga A, Miki K, Yonenobu K, Ochi T and Noguchi K. Activating transcription factor 3 (ATF3) induction by axotomy in sensory and motoneurons: A novel neuronal marker of nerve injury. Mol Cell Neurosci. 2000;15:170–82.

44. Taylor BK, Roderick RE, St Lezin E and Basbaum AI. Hypoalgesia and hyperalgesia with inherited hypertension in the rat. Am J Physiol Regul Integr Comp Physiol. 2001;280:R345–54.

45. Garsik JT, Low WC and Whitehorn D. Differences in transmission through the dorsal column nuclei in spontaneously hypertensive and Wistar Kyoto rats. Brain Res. 1983;271:188–92.

